# A novel method for harmonization of PET image spatial resolution without phantoms

**DOI:** 10.1101/2024.09.30.614929

**Authors:** Felix Carbonell, Alex P. Zijdenbos, Evan Hempel, Mihály Hajós, Barry J. Bedell, Alzheimer’s Disease Neuroimaging Initiative

**Affiliations:** Biospective Inc., 1255 Peel Street, Suite 560, Montreal, QC, H3B 2T9, Canada; Cognito Therapeutics, 1218 Massachusetts Ave., Suite 200, Cambridge, MA 02138, USA

**Keywords:** Spatial resolution, PET, Hoffman phantom, Alzheimer’s disease, β-amyloid, FDG, tau, Fourier transform, logarithmic intensity plots, FWHM

## Abstract

The most common approach for estimating the spatial resolution of PET images in multi-center studies typically uses Hoffman phantom data as a surrogate. Specifically, the phantom-based matching resolution approach assumes that scanned phantom PET images are well approximated by a ground truth, noise-free digital phantom convolved with a Gaussian kernel of unknown size. The size of the kernel is then estimated by an exhaustive search on the amount of blurring needed to match the smoothed digital phantom to a particular scanned phantom image. Unfortunately, Hoffman phantom images may not always be readily available, and phantom-based approaches may yield sub-optimal results. We propose a new, computational approach that allows estimation of spatial resolution directly from the PET image itself. We generalized the so-called logarithmic intensity plots method to the 3D case to perform a spatial resolution estimation in both axial and in-plane directions of the PET images. The proposed approach was applied to two different cohorts. The first cohort consisted of [18F]florbetapir amyloid PET images and matching phantoms coming from a Phase II clinical trial and includes different scanner models and/or orientation and grid reconstructions. The second cohort included β-amyloid, FDG and tau PET images from the Alzheimer’s Disease Neuroimaging Initiative (ADNI) study. We obtained in-plane and axial resolution estimators that vary between 3.5 mm and 8.5 mm for both PET and matching phantom images. In both cases, we obtained small across-subject variability in groups of images sharing the same PET scanner model and reconstruction parameters. For human PET images, we also obtained a strong cross-tracer and longitudinal consistency in the spatial resolution estimators. Our novel approach does not only eliminate the need for surrogate brain phantom data, but also provides a general framework that can be applied to a wide range of tracers and other image modalities, such as SPECT.

## Introduction

During the last decade, the fast development of positron emission tomography (PET) radiotracers with high specificity for β-amyloid plaques and aggregated tau in neurofibrillary tangles has represented a paradigm shift in Alzheimer’s disease (AD) research (Clark et al., 2011; Fleisher et al., 2011; Johnson et al., 2016). Indeed, the approval of several fluorine-18-labeled tracers by regulatory authorities has facilitated the use of Amyloid and Tau PET imaging as part of the clinical diagnostic work-up of patients with cognitive impairment. More importantly, PET imaging studies have accelerated our understanding of the underlying molecular mechanism driving the progression of AD and its relationship with the cognitive decline of AD patients. For instance, the amount of evidence accumulated with PET imaging suggests that β-amyloid may be the driving toxic force of tau pathology and cognitive decline observed in patients with AD (Hanseeuw et al., 2019; Wang et al., 2016). More recently, PET imaging biomarkers have confirmed the presence of AD pathology in clinical trial participants (Cummings et al., 2021, 2022). Thus, PET imaging biomarkers are becoming key players in their role of auxiliary outcomes in clinical trials and the basis for accelerated approval of recent anti-amyloid drugs for AD treatments (Cummings et al., 2021, 2022; Tolar et al., 2020). Despite slow progress in developing successful AD modifying therapies (Tolar et al., 2020), a lesson learned is the necessity to continue AD research and AD drug development within more extended scenarios of PET imaging in multi-center clinical trials.

A critical step in multi-center studies is, however, uniformization of the PET imaging biomarkers in order to allow for a fair statistical between-groups comparison by reducing the bias introduced by inter-site imaging protocols and inter-scanner variability (Joshi et al., 2009). With this aim, Joshi et al. (Joshi et al., 2009) pioneered the use of Hoffman phantoms (Hoffman et al., 1990) to reduce 1) systematic inter-scanner variability due to differences in scanner resolution, and 2) attenuation and scatter correction errors. The high frequency correction due to variability in scanner resolution (Joshi et al., 2009) was aimed at bringing Hoffman phantom images from different scanner models to a common spatial resolution. In a first step, that common minimum resolution was determined by estimating the spatial resolution of each scanner model using the phantom scans. For that purpose, an incremental Full Width at Half Maximum (FWHM) Gaussian kernel library of smoothed versions of a digital phantom image (Harrison et al., 2013) was created and then matched to each of the phantom scans using the Euclidean distance as a metric of proximity between images. Once that common target resolution was determined, a second step we necessary to estimate the amount of blurring needed to bring each phantom image to that coarsest resolution among scanner models. Similar to the first step, the amount of required blurring was estimated by creating a library of smoothed images for each phantom and matching it to blurred digital phantom. Despite the Hoffman-based approach having resulted in a useful tool for large studies like ADNI, it might not be effective for more complex settings of clinical trials. First, as pointed out early by Klein et al. (Klein et al., 2014), the preparation and analysis of the Hoffman phantoms must be carried out with extreme care to avoid non-uniformities in the phantom radiotracer concentration of the fluorine-18 solution and the subsequent bias in the spatial resolution estimation. More importantly, the employment of Hoffman phantom images poses significant logistic challenges in large and time-extended clinical trials. First, Hoffman phantom images may not always be readily available for all scanner models involved in large clinical trials. Second, one would need constant monitoring of scanner updates or replacements, including changes to the scanning protocols and image reconstruction guidelines (Schmidt et al., 2015). Additionally, the basic underlying assumption that actual human brain PET images inherit the same spatial resolution of Hoffman phantom images scanned on identical scanner models might be difficult to justify in some cases, particularly for radiotracers other than FDG.

Rather than using Hoffman phantom images as a surrogate, we introduce an approach that estimates the spatial resolution from the actual human brain PET images. The idea of estimating spatial resolution directly from real images is not a new concept and has been widely studied in the X-ray imaging and microscopy imaging fields (Brostrom & Molhave, 2022; Liao & Frank, 2010; Mizutani et al., 2010, 2016, 2017; Modregger et al., 2007; Saiga et al., 2018; Vila-Comamala et al., 2012). What is common to most of the previous methods is that spatial resolution of images is better estimated by employing Fourier analysis techniques. In particular, Mizutani *et al*. (Mizutani et al., 2016) introduced a relatively simple method for estimating the spatial resolution of 2D images from the so-called logarithmic intensity plots. A logarithm intensity plot is a scatter plot of the logarithm of the square norm of the image Fourier transform against the square distance from the origin in the frequency domain. In such a plot, the FWHM of the image is determined by a linear regression of the left end of the plot. In this manuscript, we extended the logarithmic intensity plots to the 3D case by allowing the simultaneous estimation of the FWHM in the in-plane and axial directions, as a particular case of our estimation process. Similar to the 2D case of (Mizutani et al., 2016), our main underlying assumption is that the scanned image is a lower resolution representation of an unknown higher resolution image that has been spatially convolved with Gaussian kernel of unknown size. In this sense, our estimation process follows the same basic mathematical assumptions of a Blind Deconvolution problem. The main difference is that we are not interested in recovering the underlying higher resolution image, but instead aim to estimate the size of the unknown Gaussian kernel.

Our <u>S</u>ingle <u>P</u>ET Image <u>T</u>o <u>F</u>ind Intrinsic <u>R</u>esolution <u>E</u>stimation (SPITFIRE) approach does not require any surrogate Hoffman phantom image for estimating the spatial resolution of real human brain PET images. It can be used to estimate not only the spatial resolution of real human brain PET images coming from any radiotracer and reconstructed with any grid shape and orientation, but to estimate the spatial resolution of any 2D/3D image. The performance of the SPITFIRE approach has been assessed using two different cohorts of PET images. The first cohort consists of [18F]florbetapir amyloid PET images and matching phantoms from a multi-center clinical trial that used different scanner models in its protocol. The second cohort includes amyloid, FDG, and Tau PET images from the Alzheimer’s Disease Neuroimaging Initiative (ADNI) study.

## Materials and Methods

### Subjects and Image Acquisition

The first cohort of PET images and phantom images (Cohort-1) used in the article were obtained from a multi-center study of Cognito Therapeutics’ OVERTURE clinical trial (NCT03556280) (Da et al., 2024). The Clinical Investigation plan for was developed in accordance with the requirements set forth in the United States Code of Federal Regulation, 21 CFR 812 Investigational Device Exemptions, ISO 14155:2011 Clinical Investigations of Medical Devices for Human Subjects, the Medical Device Directive 93/42/EEC of the European Union, and the Declaration of Helsinki by the World Health Organization (as amended in 2008). All individuals enrolled in the OVERTURE trial were more than 50 years old at the time of consent, had a Mini-Mental State Exam (MMSE) score ranging from 14-26, inclusive and had a diagnosis of a clinical syndrome of cognitive impairment consistent with mild to moderate AD per National Institute on Aging-Alzheimer’s Association (NIA-AA) diagnostic criteria. The OVERTURE cohort consists of n=311 human brain amyloid PET images coming from different time visits of 92 individuals and n=15 matching phantoms. The n=311 human brain amyloid ([18F]florbetapir) PET images come from 5 different combinations of site ID/manufacturer/scanner model as shown in Table 1.

**Table 1.** Distribution of Cohort-1 amyloid PET data.

| Site ID | Manufacturer | Scanner Model | N | Image Size | Voxel Size |
| --- | --- | --- | --- | --- | --- |
| A | GE MEDICAL SYSTEMS | Discovery STE | 196 | 128x128x47 | 2x2x3.27 |
| B | GE MEDICAL SYSTEMS | Discovery ST | 41 | 128x128x47 | 2x2x3.27 |
| C | SIEMENS | Biograph Horizon | 25 | 256x256x81 | 1.015x1.015x2.025 |
| D | SIEMENS | Biograph6 TruePoint | 25 | 336x336x83 | 1.018x1.018x2.0 |
| E | SIEMENS | Biograph20_mCT | 24 | 400x400x81 | 1.018x1.018x2.025 |

There was a total of n=15 phantoms matching the n=311 human brain PET images according to their site ID, manufacturer and scanner model and distributed as A (n=5), B (n=3), C (n=2), D (n=1) and E (n=4).

The second cohort of PET images used in the preparation of this article were obtained from the Alzheimer’s Disease Neuroimaging Initiative (ADNI, http://adni.loni.usc.edu). The ADNI project was launched in 2003 by the National Institute on Aging (NIA), the National Institute of Biomedical Imaging and Bioengineering (NIBIB), the Food and Drug Administration (FDA), private pharmaceutical companies, and non-profit organizations, as a $60 million, 5-year public private partnership, which has since been extended. ADNI is the result of efforts of many co-investigators from a broad range of academic institutions and private corporations, and subjects have been recruited from over 55 sites across the U.S. and Canada. To date, the ADNI, ADNI-GO, ADNI-2 and ADNI-3 protocols have recruited over 1,500 adults, ages 55 to 90, to participate in the research, consisting of cognitively normal (CN) older individuals, people with early or late MCI, and people with early AD. For up-to-date information, see www.adni-info.org.

The ADNI sample used in this study consisted of 3,419 PET images, including [18F]Florbetapir PET (n=1706), [18F]FDG PET (n=999), and [18F]Flortaucipir PET (n=714). A detailed description of the ADNI MRI and PET image acquisition protocols can be found at http://adni.loni.usc.edu/methods. ADNI studies are conducted in accordance with the Good Clinical Practice guidelines, the Declaration of Helsinki, and U.S. 21 CFR Part 50 (Protection of Human Subjects), and Part 56 (Institutional Review Boards), where informed written consent was obtained from all participants at each site.

### Image Processing

PET images were processed using the PIANO™ software package (Biospective Inc., Montreal, Canada). The human brain PET images were submitted to a frame-to-frame linear motion correction followed by averaging of dynamic frames into a static image. The resulting static PET images were then submitted to our proposed SPITFIRE approach for resolution estimation. To assess the effect of the spatial resolution estimation in the final processed PET images, a common target resolution of 8 mm was selected, and a specific spatial blurring was applied to each of the human brain PET images. The resulting smoothed PET volumes were linearly registered to the subject’s T1-weighted MRI, where the T1-weighted MRI volumes previously underwent image non-uniformity correction using the N3 algorithm [22], brain masking, linear spatial normalization utilizing a 9-parameter affine transformation, and nonlinear spatial normalization to Montreal Neurological Institute (MNI) reference space using a customized, anatomical MRI template. An anatomical atlas has been constructed and it is composed of several Regions of Interest (ROIs) that has been defined in the anatomical MRI template. ROIs are mapped onto the PET scan and used to collect ROI-based SUVR standardized uptake value ratio (SUVR) measures for any reference region using automated atlas-based parcellation in stereotaxic space. The PET processing pipeline excludes voxels located outside of the PET field-of-view (FOV) when computing average ROI values. Thus, the maps of the SUVR PET images were generated from [18F]florbetapir, [18F]FDG, and [18F]flortaucipir PET using full cerebellum, pons, and full cerebellum as reference regions, respectively.

### SPITFIRE

Our proposed approach is based on the generalization of the 2D logarithmic intensity plots in the Fourier domain (Mizutani et al., 2016) to the 3D case. The basic element in the original 2D logarithmic intensity plots approach is to plot the logarithm of the square norm of the image Fourier transform against the square distance from the origin in the frequency domain. Once in this plot, the unknown width of the Gaussian kernel is determined by a linear regression of the left side of the plot (i.e., smallest square frequency values).

Our generalization does not only apply to the estimation of isotropic resolution in 3D, but also allows to consider a different in-plane and axial resolution spatial estimations. The new SPITFIRE approach also assumes that any (noisy) scanned PET image is well approximated by an unknown higher resolution image convolved with a Gaussian kernel of unknown size. The sizes of the kernel for the in-plane and axial directions are then estimated by multiple linear regression involving the observed image power spectrum. Specifically, let’s denote by *Y(v)* the 3D PET image at voxel v. Then, our assumptions can be written as *Y*(*v*) = *X*(*v*)⊗*k*(*σ*_x_, *σ*_y_, *σ*_z_), where *X(v)* is the unknown, higher resolution image, *k*(*σ*_x_, *σ*_y_, *σ*_z_) is a 3D Gaussian kernel determined by the kernel widths *σ*_x_, *σ*_y_ and *σ*_z_ in each of the three spatial directions and ⊗ denotes the 3D convolution operator. Using the fact that fast Fourier transform (FFT) of a Gaussian kernel also produces a Gaussian kernel in the 3D frequency domain, we can obtain 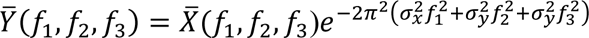, where 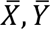 are 3D images in the Fourier domain defined by the three frequencies *f*_1_, *f*_2_, *f*_3_. Thus, similarly to (Mizutani et al., 2016), taking the power spectrum and logarithm on both sides of the expression above yields a multiple regression problem in the unknown variables *σ*_x_, *σ*_y_, *σ*_z_. By using the additional assumption of *σ*_x_ = *σ*_y_ = *σ*_xy_, we can determine the in-plane and axial image resolutions as *FWHM*_xy_ = 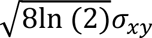 and *FWHM*_z_ = 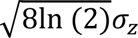, respectively.

As suggested by Mizutani *et al*. (Mizutani et al., 2016), in order to avoid undesired noise effects during the linear regression step, we restricted the regression to those values corresponding to the top 25% for each of the frequency values *f*_1_, *f*_2_, *f*_3_. We also eliminated those values containing the zero frequency from the regression step.

Finally, the amount of blurring needed in order to achieve a desired target FWHM value can be derived from the properties of Gaussian kernels according to the explicit analytical formula FWHM^2^_to-target_ = FWHM^2^_target_ - FWHM^2^_estimated_, where *FWHM*_estimated_ denotes the spatial resolution estimated from the actual image, namely by the SPITFIRE approach.

## Results

The SPITFIRE approach was applied to each of the n=311 human brain PET images and n=15 phantom images from Cohort-1. For presentation purposes, the images were arranged according to their corresponding site ID, manufacturer, scanner model, and reconstruction parameters (e.g., image dimensions and voxel size). The mean and standard deviation values of axial and in-plane FWHMs for both human brain PET data and phantoms are presented in Table 2.

**Table 2.**
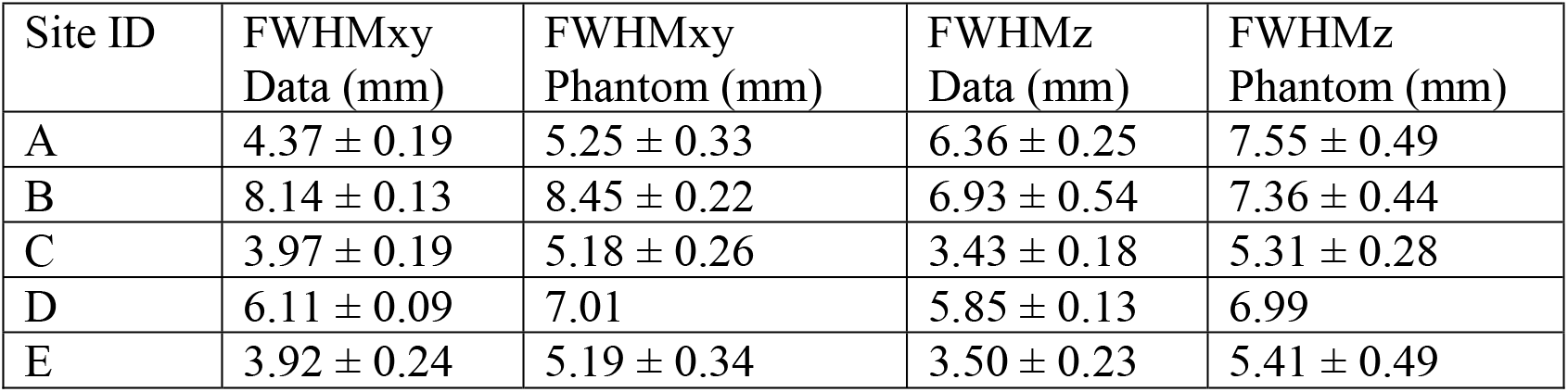
Resolution estimator of Cohort-1 PET and phantom data.

| Site ID | FWHM<sub>xy</sub><br>Data (mm) | FWHM<sub>xy</sub><br>Phantom (mm) | FWHM<sub>z</sub><br>Data (mm) | FWHM<sub>z</sub><br>Phantom (mm) |
| --- | --- | --- | --- | --- |
| A | $4.37 \pm 0.19$ | $5.25 \pm 0.33$ | $6.36 \pm 0.25$ | $7.55 \pm 0.49$ |
| B | $8.14 \pm 0.13$ | $8.45 \pm 0.22$ | $6.93 \pm 0.54$ | $7.36 \pm 0.44$ |
| C | $3.97 \pm 0.19$ | $5.18 \pm 0.26$ | $3.43 \pm 0.18$ | $5.31 \pm 0.28$ |
| D | $6.11 \pm 0.09$ | 7.01 | $5.85 \pm 0.13$ | 6.99 |
| E | $3.92 \pm 0.24$ | $5.19 \pm 0.34$ | $3.50 \pm 0.23$ | $5.41 \pm 0.49$ |

Note that in most cases, the FWHM estimation of the human brain data is within the tolerance range (less than the voxel size) of the matching phantom FWHM estimation. Exceptions are the C and the E site IDs where the mean FWHMxy estimation is a bit smaller than one voxel size of the mean FWHMxy of those matching phantoms. However, even in those cases, the corresponding confidence interval (i.e., accounting for the standard deviation as well) overlap, meaning that those mean values could be considered statistically close each other.

The longitudinal stability of the FWHM estimation in human brain images was evaluated by computing the maximum absolute difference of the resolution estimation across all time visits, for each individual subject. Figure 1 shows this maximum difference metric for all subjects, where each color represents a unique site ID/manufacturer/scanner model (ordered by alphabetically by site ID). In this figure, the maximum difference of 2.20 mm corresponds to a subject coming from the site B and producing an FWHMz estimation of 6.78 mm, 7.96 mm, and 5.76 mm for the baseline and two follow-up visits. Note, however, that the voxel size in the axial direction for the site B was 3.27 mm, which means that the maximum longitudinal variability is less than a voxel size.

**Figure 1.**
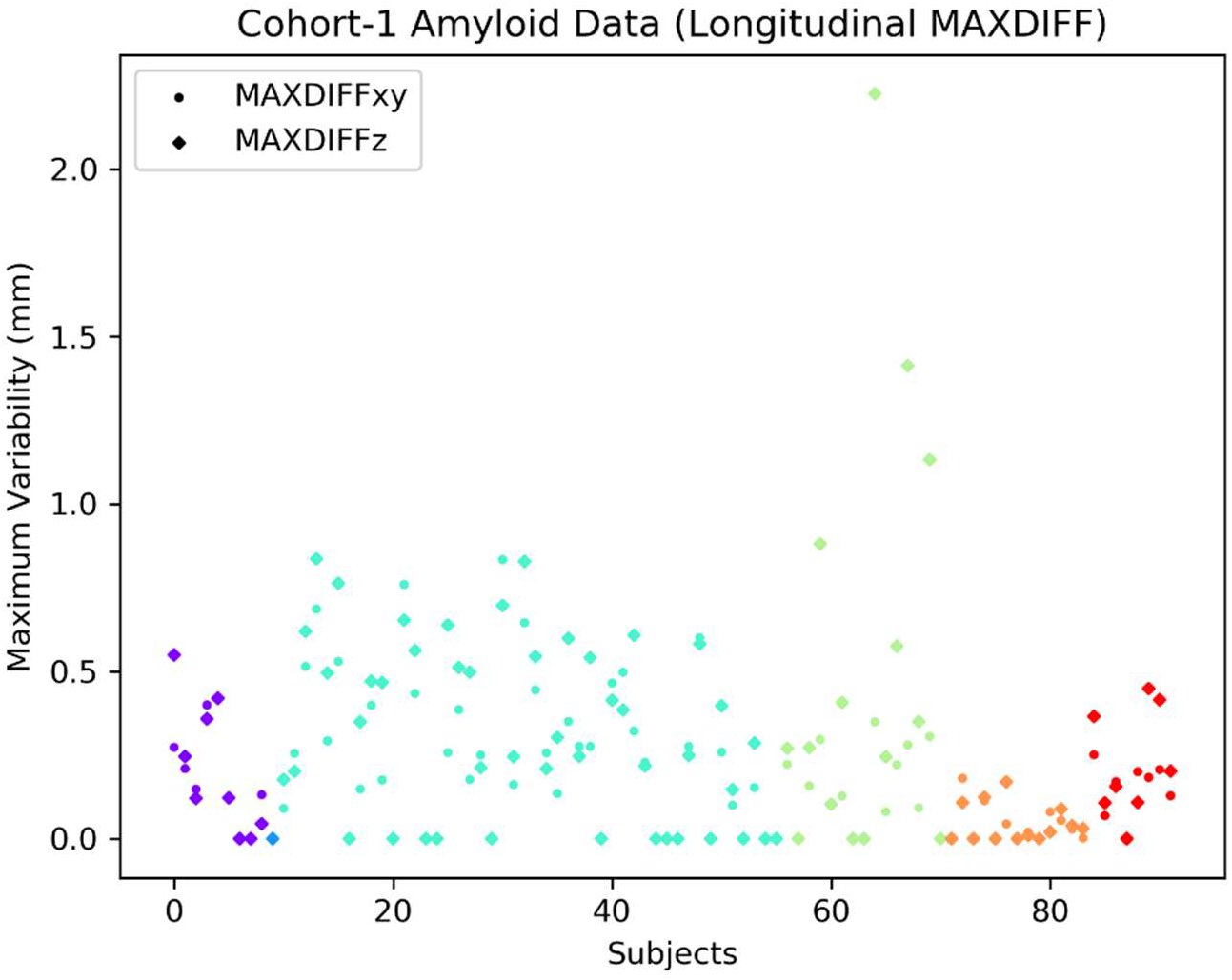
Longitudinal stability of FWHM estimation in Cohort-1 Amyloid data.

The SUVR comparison of data processed with Hoffman phantom kernels and the SPITFIRE approach is presented in Figure 2 for several ROIs: Anterior Cingulate Cortex, Occipital Cortex, Deep White Matter, and Pons, where the SUVR was computed using the cerebellum as a reference region and each color represents a unique site ID/manufacturer/scanner model as in Table 1.

**Figure 2.**
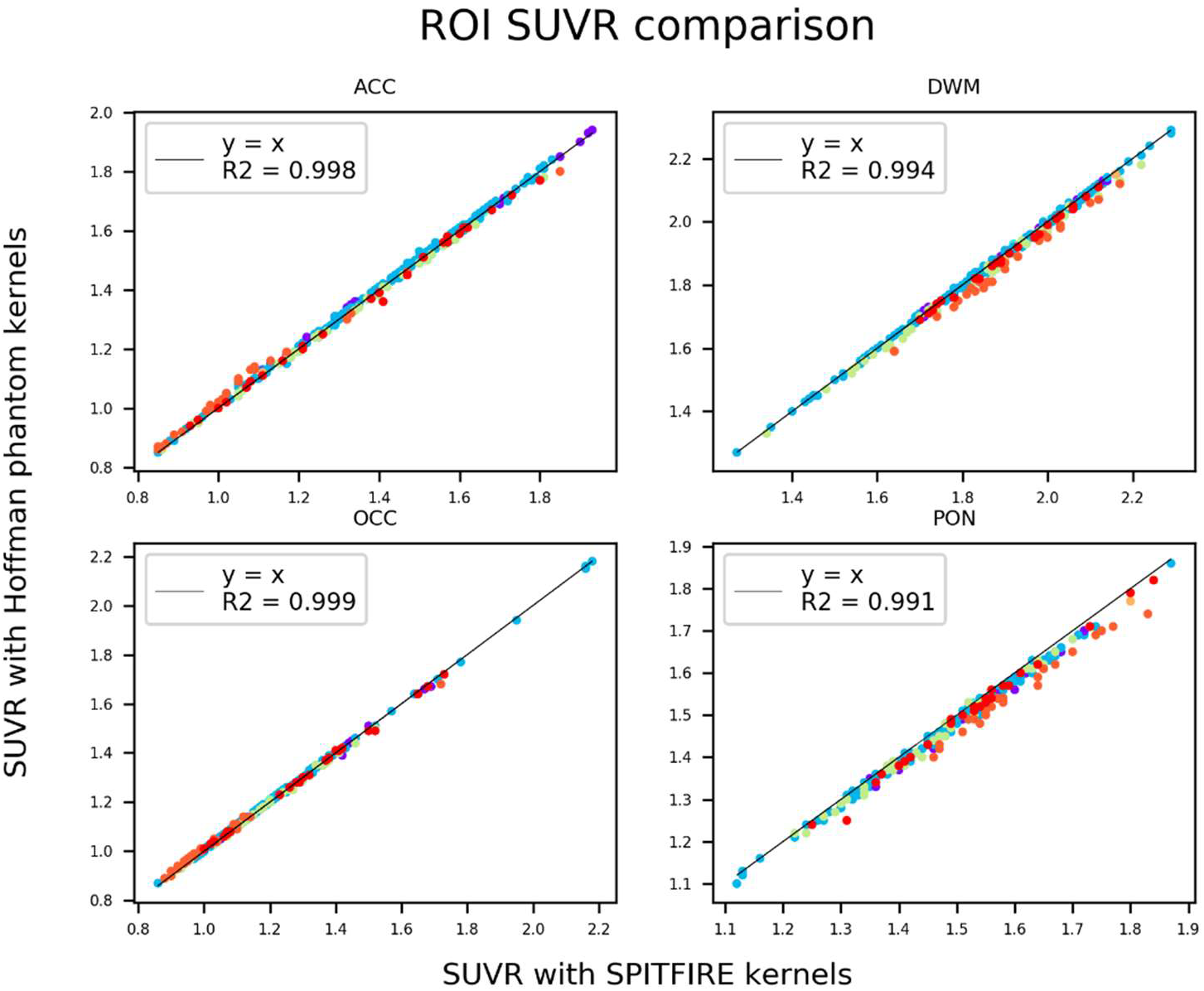
SUVR comparison for ROI SUVR Cohort-1 Amyloid data.

Notice that the SUVR values obtained with both approaches are extremely similar (r > 0.99) with only small differences for the Pons ROI probably due to the very small size of this region.

Although only 4 ROIs were presented in the previous figure, the correlation coefficient was also computed for the following ROIs: Frontal cortex, Lateral temporal cortex, Medial temporal cortex, Occipital pole, Parietal cortex, Posterior cingulate cortex, Precuneus cortex, Sensorimotor cortex, Striatum, Thalamus, and Ventral Striatum. In all cases, the correlation coefficient was also greater than 0.99.

The SPITFIRE approach was also applied to each of the PET images in the ADNI cohort. As in the case of Cohort-1, the images were arranged according to their site ID, scanner manufacturer/model, and reconstruction parameters. Supplementary Table 1 shows the SPITFIRE resolution estimator of ADNI amyloid data for those sites with more than 15 images coming from the same scanner manufacturer/model and sharing the same reconstruction parameters. Similarly, Supplementary Table 2 and Supplementary Table 3 show the resolution estimator of ADNI FDG and Tau data, respectively.

Figure 3 shows the boxplots distribution corresponding to the FWHM estimation in both, axial and in-plane directions for several sites in the ADNI cohort.

**Figure 3.**
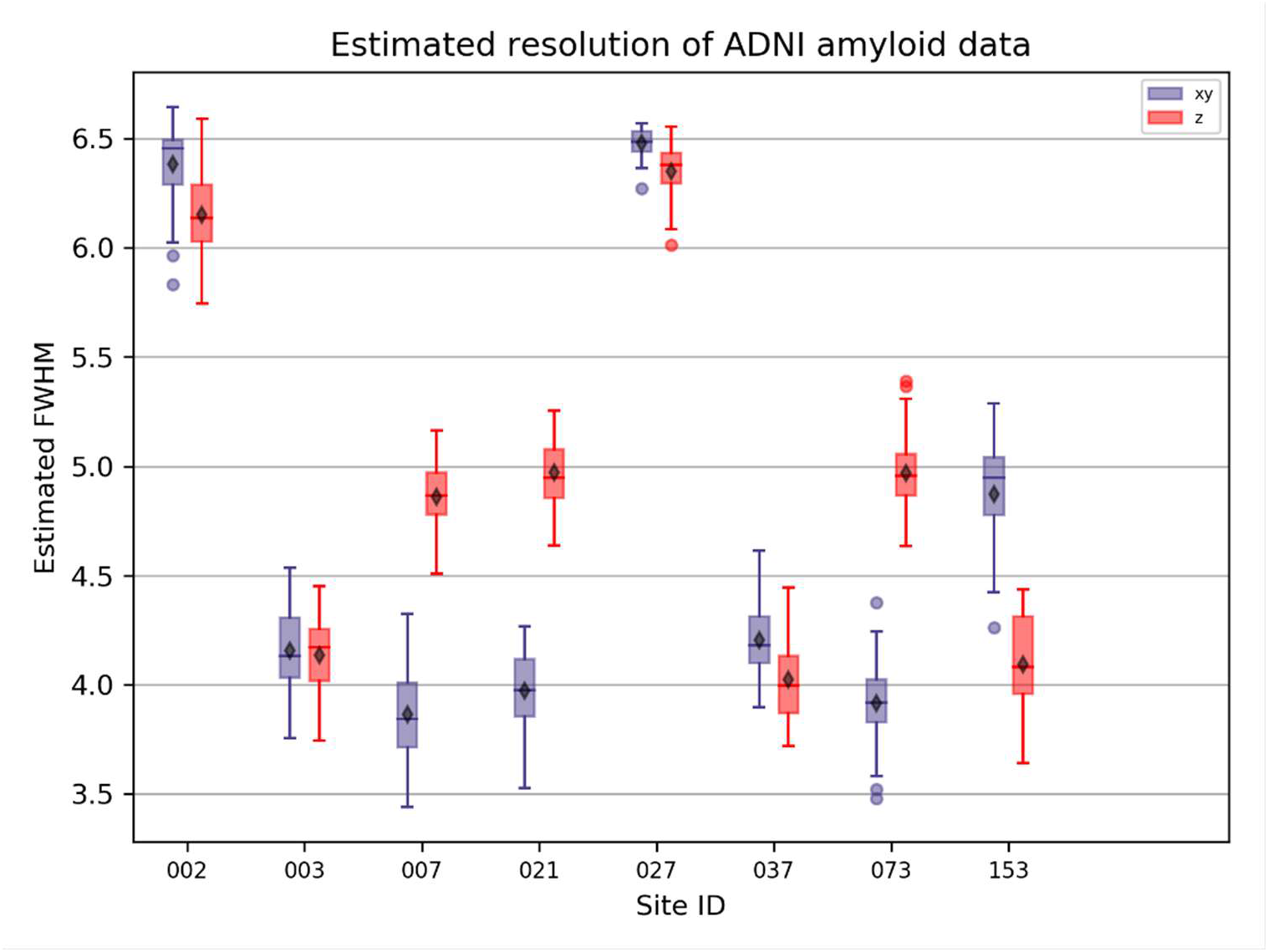
FWHM estimation of source ADNI Amyloid data.

Notice that, in order to cover a large range of different site/manufacturer/scanner models, only a few sites are shown in the previous figure. The full list of SPITFIRE estimators can be seen in Supplementary Table 1. Notice also that, in the previous figure, the mean value of the in-plane SPITFIRE estimators (i.e., FWHMxy) tends to be very similar or less than the axial estimator FWHMz for most of the site IDs. The most marked case where the mean FWHMxy is evidently larger than FWHMz is the site ID labelled as 153, which corresponds to a Biograph20_mCT/SIEMENS model with image shapes of 256 x 256 x 81 voxels and voxel size of 1.6 x 1.6 x 2.02 mm. The maximum absolute difference of the SPITFIRE estimator was computed for all individual subjects in the ADNI cohort and the three PET tracers. Figure 4 shows the boxplots for this maximum difference metric across all individual subjects with more than one time point visit.

**Figure 4.**
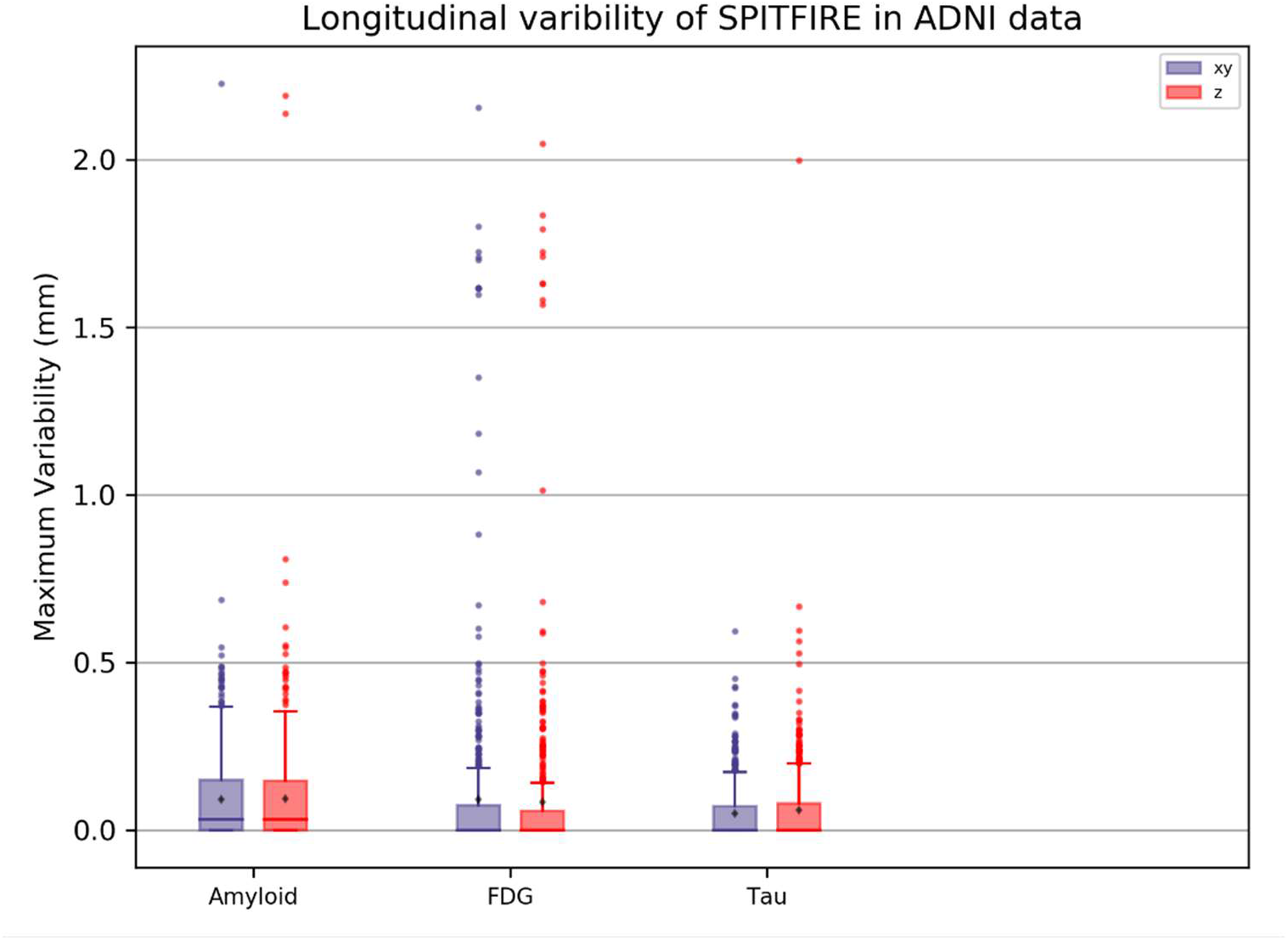
Longitudinal stability of SPITFIRE estimator in ADNI cohort.

Careful verification of the data shown in the previous figure revealed that, in all cases, the maximum difference among time visits resolution estimation was less than the corresponding voxel size.

To assess the consistency of the SPITFIRE estimator across tracers, we selected subjects with pairs of PET images with different tracers, but scanned no more than 3 months apart. Thus, Figure 5 shows scatter plots of the resolution estimation for each pair of tracers.

**Figure 5.**
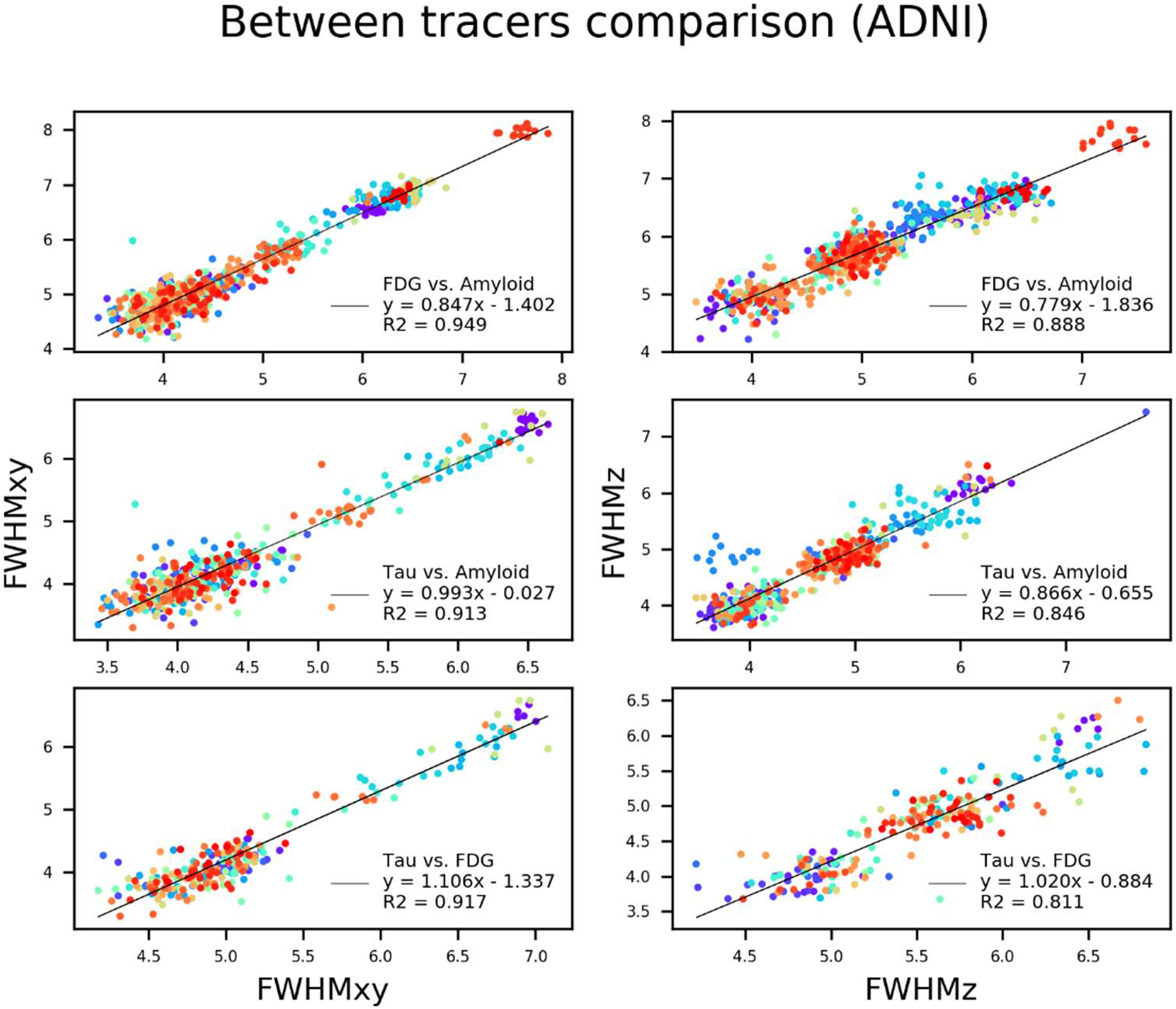
Between-tracer comparison of the SPITFIRE estimator in ADNI cohort.

Notice that in each case the correlation between tracers is slightly greater for the in-plane resolution as compared to the axial resolution, perhaps driven by the typically greater voxel size in the axial direction.

## Discussion

By using the SPITFIRE approach, we obtained resolution estimator values that showed very small (less than a voxel size) across-subject variability when the subjects were aggregated by site/manufacturer/scanner model and reconstruction parameters. More importantly, the estimation of resolution of actual human PET images closely matches that of the corresponding phantom data from the same scanner model and identical reconstruction parameters. Thus, in contrast to the Hoffman-based approach (Joshi et al., 2009) that assumes a common spatial resolution value for images acquired using the same scanner model, our SPITFIRE approach can produce scan-specific spatial resolution estimations. Hence, the SPITFIRE technique is not only able to capture systematic differences in spatial resolution related to scanner hardware and reconstruction software but also account for inter-subject variability due to anatomical and functional differences.

There are also key differences between the Hoffman-based and the SPITFIRE approaches that are worth mentioning. A main element is the initial preprocessing of phantom scans prior estimation of spatial resolution. As it was mentioned in (Joshi et al., 2009), the image volume of each phantom scan is initially registered to the digital Hoffman brain phantom to produce a common orientation and image shape for all phantom scans. Thus, the estimated resolution of the registered Hoffman phantom differs from the original resolution of the Hoffman phantom due to the spatial transformation (e.g., resampling) required to align the original phantom scan to the digital Hoffman phantom volume (Worsley et al., 1992). Indeed, as a simple example, Worsley et al. (Worsley et al., 1992) provided explicit expressions for the change in spatial smoothness resulting from resampling (i.e., by linear interpolation) of PET images in the axial direction. Since the Hoffman-based approach for human PET images is based on the assumption that human data inherit the same spatial resolution of the matching phantom, it is likely to conclude that such assumption is only valid when the actual human PET image is registered to the same grid and orientation of the digital Hoffman phantom. Otherwise, a correction on spatial resolution estimation like the one presented in (Worsley et al., 1992) would be in order. On the other hand, compared to the Hoffman-based approach of (Joshi et al., 2009), SPITFIRE can be applied to images in any orientation and shape, regardless of if they were previously resampled. Indeed, SPITFIRE would account for the effective spatial resolution and no extra correction step would be required. In fact, the application of SPITFIRE in this manuscript only assumes that actual human multi-frame PET volumes have been submitted to an initial motion correction step before transformation to a static 3D image. This way, SPITFIRE also accounts for any change in smoothness introduced by the motion correction step, as compared to the original scanned image. Another aspect that has only rarely been discussed since the inception of the Hoffman-based approach is its validity for PET tracers other than FDG. While the scanned Hoffman phantoms in (Joshi et al., 2009) were filled with a fluorine-18 solution, their matching resolution approach was only assessed for [18F]FDG human brain PET data. To our knowledge, no subsequent publication has addressed the validity of this scanner-matching approach for other PET tracers like [18F]florbetapir or [18F]flortaucipir. Correspondingly, another important issue to address is the inter-tracer variability in subjects scanned with different PET tracers and at different time points. In this regard, our results with the SPITFIRE approach revealed that the spatial resolution estimation was highly consistent across tracers (see Figure 5), producing between-tracer correlations higher than r=0.9. We also corroborated that the SPITFIRE approach is longitudinally stable in the sense that it produced a strong consistency in resolution estimation across PET images from the same subject at different time points. This longitudinal consistency was assessed by computing the maximum absolute difference among all pairs of time points estimators and confirming that they were in the order of less than the corresponding voxel size. An advantage of the SPITFIRE approach over the Hoffman-based procedure of (Joshi et al., 2009) is related to the computation of blurring kernels to achieve a desired common resolution among different scanners. While the Hoffman-based approach of (Joshi et al., 2009) requires a library of incremental FWHM Gaussian kernels for each phantom scan, the SPITFIRE approach relies on the well-known properties of Gaussian kernels and uses an analytical formula to achieve any desired target resolution. Indeed, even the Hoffman-based approach could benefit of this analytical formula and avoid the unnecessary step of matching the library of blurred phantoms to the blurred digital phantom.

The use of phantom scans as a surrogate for estimating the spatial resolution of actual human brain PET images comes with several limitations and pitfalls that can be avoided using SPITFIRE. As pointed out by Klein et al. (Klein et al., 2014), the preparation and analysis of 3D Hoffman phantoms for human brain PET imaging is not a trivial task. Indeed, an improper preparation of the phantoms typically produce air bubbles that create regions of hypointensity and non-uniformities in the radiotracer concentration of the fluorine-18 solution (Klein et al., 2014). Another common pitfall is that, for many scanners, Hoffman phantoms do not cover the totality of the axial field of view (Klein et al., 2014). A major limitation of that the Hoffman-based approach forces an initial pre-blurring of all the PET images to a common lower resolution target. Thus, even high-resolution images are spatially degraded at the very beginning of the pre-processing pipeline for the sole purposes of having a final common resolution that is only necessary for statistical purposes. Thus, intermediate preprocessing steps, such as PET-MRI co-registration, may be affected by this initial pre-blurring of originally high-resolution PET images. In fact, it is likely that the final processed images do not bear a common spatial resolution after all. Indeed, having a common initial resolution of images from different grids and orientations does not guarantee that the final processed images bear the common target resolution since the pre-processing typically involves several re-sampling steps, as well as linear and non-linear image registration and normalization steps that certainly modifies (Worsley et al., 1992) the effective spatial resolution. In this context, the SPITFIRE approach can be applied at any stage of the pre-processing pipeline. Thus, blurring to a common target resolution can be introduced as the final step of the PET pre-processing pipeline as a necessary statistical correction, more in correspondence with the classical PET studies involving data coming from a single center and a single scanner.

The applicability of any pre-blurring approach in multi-center clinical trial faces a common limitation, namely, the *a priori* determination of the final target resolution. We then envision using our SPITFIRE approach on a few initial images scanned at the beginning of the study in order to determine each scanner resolution and propose a rather liberal target FWHM greater than the coarsest resolution among the initial images. Since the resolution will be initially estimated from each image data at scan level, this approach is considerably less sensitive to scanner changes than in most of the multi-center clinical trials.

Last but not least, besides the methodological advantages described above, the use of SPITFIRE obviates the need to collect Hoffman phantom scans in clinical studies, which is not only costly, but logistically challenging. For example, changes to scanner hardware/software, that would require re-estimating the scanner resolution using another Hoffman phantom scan, may not be known in a timely fashion, hampering the association of all subject scans with a representative Hoffman phantom scan. Especially in large, multi-center studies, the effort required to ensure that Hoffman scans are acquired when needed, analyze them, and associate the resulting resolution estimates with the appropriate subject scans, can be substantial.

## Conclusions

We developed the SPITFIRE approach for estimating the spatial resolution of PET images using a 3-dimensional generalization of logarithmic intensity plots. Our analysis of actual human brain PET data and accompanying phantom images revealed spatial resolution values with very low variability among images coming from the same scanner model and reconstruction parameters. For all cases, the variability across images aggregated in the same group according to their scanner model was of less than the corresponding voxel size. The SPITFIRE estimation also showed a very strong longitudinal stability from PET images coming from the same subject at different scanning time points. Additionally, the SPITFIRE produced a very strong inter-tracer consistency on those subjects scanned at close time points with different radiotracers. To our knowledge, this study is the first to provide a clear methodology to estimate the spatial resolution of real PET images without the need to rely on accompanying Hoffman phantom images and the subsequent tedious and biased image-to-phantom matching process. Our results showed that the proposed approach can be readily used in large clinical trial environments, considerably decreasing the cost and operational effort of acquiring Hoffman phantoms, especially in multi-center studies.

## Acknowledgments

Data collection and sharing for this project was funded by the Alzheimer’s Disease Neuroimaging Initiative (ADNI) (National Institutes of Health Grant U01 AG024904) and DOD ADNI (Department of Defense award number W81XWH-12-2-0012). ADNI is funded by the National Institute on Aging, the National Institute of Biomedical Imaging and Bioengineering, and through generous contributions from the following: AbbVie, Alzheimer’s Association; Alzheimer’s Drug Discovery Foundation; Araclon Biotech; BioClinica, Inc.; Biogen; Bristol-Myers Squibb Company; CereSpir, Inc.; Cogstate; Eisai Inc.; Elan Pharmaceuticals, Inc.; Eli Lilly and Company; EuroImmun; F. Hoffmann-La Roche Ltd and its affiliated company Genentech, Inc.; Fujirebio; GE Healthcare; IXICO Ltd.; Janssen Alzheimer Immunotherapy Research & Development, LLC.; Johnson & Johnson Pharmaceutical Research & Development LLC.; Lumosity; Lundbeck; Merck & Co., Inc.; Meso Scale Diagnostics, LLC.; NeuroRx Research; Neurotrack Technologies; Novartis Pharmaceuticals Corporation; Pfizer Inc.; Piramal Imaging; Servier; Takeda Pharmaceutical Company; and Transition Therapeutics. The Canadian Institutes of Health Research is providing funds to support ADNI clinical sites in Canada. Private sector contributions are facilitated by the Foundation for the National Institutes of Health (www.fnih.org). The grantee organization is the Northern California Institute for Research and Education, and the study is coordinated by the Alzheimer’s Therapeutic Research Institute at the University of Southern California. ADNI data are disseminated by the Laboratory for Neuro Imaging at the University of Southern California.

## Author contributions

FC and BJB designed research; FC, APZ, and BJB performed research; FC and BJB analyzed the data and wrote the paper.

## Compliance with ethical standards

### Ethical approval

All procedures performed in studies involving human participants were in accordance with the ethical standards of the institutional and/or national research committee and with the 1964 Helsinki declaration and its later amendments or comparable ethical standards.

### Informed consent

Informed consent was obtained from all individual participants included in the study.

