## Supplementary Table 1 for "A novel method for harmonization of PET image spatial resolution without phantoms"

| **Site** | **Model** | **Manufacturer** | **FWHMxy Mean** | **FWHMxy Std** | **FWHMz Mean** | **FWHMz Std** | **N** |
| --- | --- | --- | --- | --- | --- | --- | --- |
| 002 | GEMINI TF TOF 16 | Philips Medical Systems | 6.38 | 0.17 | 6.15 | 0.19 | 84 |
| 073 | Discovery STE | GE MEDICAL SYSTEMS | 3.92 | 0.17 | 4.97 | 0.15 | 74 |
| 018 | Advance | GEMS | 4.37 | 0.28 | 5.5 | 0.16 | 49 |
| 033 | Discovery ST | GE MEDICAL SYSTEMS | 5.28 | 0.16 | 4.8 | 0.14 | 47 |
| 126 | Discovery STE | GE MEDICAL SYSTEMS | 3.96 | 0.18 | 5.03 | 0.17 | 46 |
| 003 | 1094 | SIEMENS | 4.16 | 0.20 | 4.14 | 0.17 | 43 |
| 027 | Ingenuity TF PET/CT | Philips | 6.21 | 0.12 | 5.69 | 0.21 | 41 |
| 009 | Discovery LS | GE MEDICAL SYSTEMS | 4.43 | 0.21 | 5.6 | 0.52 | 38 |
| 003 | Biograph64_TruePoint | SIEMENS | 4.32 | 0.22 | 4.01 | 0.17 | 37 |
| 016 | LSO PET/CT HI-REZ | SIEMENS | 4.25 | 0.34 | 3.84 | 0.16 | 37 |
| 098 | Discovery STE | GE MEDICAL SYSTEMS | 3.95 | 0.21 | 4.94 | 0.12 | 35 |
| 116 | Discovery ST | GE MEDICAL SYSTEMS | 4.23 | 0.17 | 4.94 | 0.15 | 34 |
| 067 | 1094 | SIEMENS | 4.12 | 0.23 | 4.21 | 0.20 | 33 |
| 007 | Discovery RX | GE MEDICAL SYSTEMS | 3.86 | 0.21 | 4.86 | 0.15 | 33 |
| 130 | Biograph 64_mCT | SIEMENS | 4.17 | 0.25 | 4.04 | 0.22 | 33 |
| 137 | Discovery STE | GE MEDICAL SYSTEMS | 4.27 | 0.22 | 5.00 | 0.12 | 32 |
| 130 | SOMATOM Definition AS_mCT | SIEMENS | 5.00 | 0.19 | 4.21 | 0.19 | 31 |
| 137 | Discovery ST | GE MEDICAL SYSTEMS | 5.16 | 0.15 | 4.74 | 0.15 | 30 |
| 027 | GEMINI TF TOF 64 | Philips Medical Systems | 6.48 | 0.07 | 6.35 | 0.13 | 29 |
| 011 | 1094 | SIEMENS | 4.15 | 0.16 | 4.25 | 0.14 | 29 |
| 037 | Biograph128_mCT | SIEMENS | 4.20 | 0.16 | 4.02 | 0.18 | 29 |
| 019 | Discovery 600 | GE MEDICAL SYSTEMS | 3.68 | 0.23 | 4.82 | 0.21 | 27 |
| 941 | Discovery ST | GE MEDICAL SYSTEMS | 4.22 | 0.17 | 4.82 | 0.14 | 24 |
| 021 | Discovery STE | GE MEDICAL SYSTEMS | 4.12 | 0.22 | 5.07 | 0.20 | 23 |
| 168 | Discovery STE | GE MEDICAL SYSTEMS | 4.16 | 0.23 | 5.00 | 0.13 | 22 |
| 941 | GEMINI TF TOF 16 | Philips Medical Systems | 6.35 | 0.07 | 6.40 | 0.13 | 22 |
| 006 | Biograph6_TruePoint | SIEMENS | 4.02 | 0.13 | 3.78 | 0.15 | 22 |
| 022 | Ingenuity TF PET/CT | Philips | 5.98 | 0.15 | 5.92 | 0.22 | 22 |
| 012 | Discovery 710 | GE MEDICAL SYSTEMS | 3.91 | 0.28 | 4.80 | 0.17 | 21 |
| 035 | Biograph64_TruePoint | SIEMENS | 4.17 | 0.15 | 4.04 | 0.13 | 21 |
| 153 | Biograph20_mCT | SIEMENS | 4.87 | 0.26 | 4.09 | 0.23 | 20 |
| 135 | SCIN | MiE | 5.33 | 0.26 | 5.06 | 0.19 | 20 |
| 021 | Discovery 690 | GE MEDICAL SYSTEMS | 3.97 | 0.18 | 4.97 | 0.16 | 19 |
| 135 | Discovery MI | GE MEDICAL SYSTEMS | 3.80 | 0.21 | 4.68 | 0.17 | 19 |
| 130 | Biograph64 | SIEMENS | 4.71 | 0.16 | 4.31 | 0.21 | 19 |
| 027 | Vereos PET/CT | Philips | 5.58 | 0.18 | 5.2 | 0.17 | 18 |
| 022 | GEMINI TF Big Bore | Philips Medical Systems | 6.30 | 0.09 | 6.32 | 0.22 | 18 |
| 035 | 1094 | SIEMENS | 4.03 | 0.14 | 4.12 | 0.09 | 18 |
| 037 | SOMATOM Definition AS_mCT | SIEMENS | 4.64 | 0.16 | 4.06 | 0.21 | 17 |
| 100 | Discovery STE | GE MEDICAL SYSTEMS | 4.12 | 0.13 | 5.00 | 0.09 | 17 |
| 141 | Guardian Body(C) | Philips Medical Systems | 7.59 | 0.14 | 7.25 | 0.2 | 17 |
| 057 | Discovery STE | GE MEDICAL SYSTEMS | 4.04 | 0.17 | 5.00 | 0.18 | 17 |
| 014 | NULL | Philips Medical Systems | 6.39 | 0.12 | 6.36 | 0.17 | 16 |
| 114 | GEMINI TF TOF 64 | Philips Medical Systems | 6.52 | 0.08 | 6.10 | 0.17 | 16 |
| 127 | Discovery MI | GE MEDICAL SYSTEMS | 4.11 | 0.25 | 4.68 | 0.22 | 15 |
| 141 | Biograph 20_mCT | SIEMENS | 4.20 | 0.22 | 4.01 | 0.19 | 15 |

**Supplementary Table 1**. SPITFIRE resolution estimator of ADNI amyloid data.
