## Supplementary Table 2 for "A novel method for harmonization of PET image spatial resolution without phantoms"

| **Site** | **Model** | **Manufacturer** | **FWHMxy Mean** | **FWHMxy Std** | **FWHMz Mean** | **FWHMz Std** | **N** |
| --- | --- | --- | --- | --- | --- | --- | --- |
| 002 | GEMINI TF TOF 16 | Philips Medical Systems | 6.75 | 0.17 | 6.62 | 0.18 | 48 |
| 073 | Discovery STE | GE MEDICAL SYSTEMS | 4.77 | 0.17 | 5.7 | 0.15 | 40 |
| 126 | Discovery STE | GE MEDICAL SYSTEMS | 4.68 | 0.15 | 5.67 | 0.15 | 31 |
| 137 | Discovery STE | GE MEDICAL SYSTEMS | 5.01 | 0.22 | 5.7 | 0.18 | 29 |
| 114 | Gemini TF(C) | Philips Medical Systems | 6.79 | 0.82 | 6.37 | 0.81 | 29 |
| 098 | Discovery STE | GE MEDICAL SYSTEMS | 4.73 | 0.18 | 5.7 | 0.13 | 27 |
| 018 | Advance | GEMS | 5.02 | 0.22 | 6.3 | 0.24 | 26 |
| 116 | Discovery ST | GE MEDICAL SYSTEMS | 4.95 | 0.26 | 5.66 | 0.22 | 25 |
| 033 | Discovery ST | GE MEDICAL SYSTEMS | 5.88 | 0.17 | 5.59 | 0.19 | 24 |
| 027 | GEMINI TF TOF 64 | Philips Medical Systems | 6.85 | 0.1 | 6.73 | 0.12 | 23 |
| 007 | Discovery RX | GE MEDICAL SYSTEMS | 4.77 | 0.16 | 5.71 | 0.15 | 21 |
| 130 | Biograph64 | SIEMENS | 5.31 | 0.21 | 4.98 | 0.22 | 21 |
| 941 | Guardian Body(C) | Philips Medical Systems | 8.01 | 0.21 | 7.74 | 0.28 | 19 |
| 130 | SOMATOM Definition AS_mCT | SIEMENS | 5.63 | 0.15 | 5.05 | 0.16 | 19 |
| 027 | Guardian Body(C) | Philips Medical Systems | 5.11 | 0.51 | 5.78 | 0.2 | 18 |
| 141 | Guardian Body(C) | Philips Medical Systems | 7.99 | 0.1 | 7.78 | 0.23 | 16 |
| 022 | GEMINI TF Big Bore | Philips Medical Systems | 6.73 | 0.09 | 6.7 | 0.17 | 16 |
| 009 | Discovery LS | GE MEDICAL SYSTEMS | 5.07 | 0.16 | 6.24 | 0.15 | 16 |
| 941 | GEMINI TF TOF 16 | Philips Medical Systems | 6.82 | 0.08 | 6.77 | 0.07 | 16 |
| 168 | Discovery STE | GE MEDICAL SYSTEMS | 4.78 | 0.24 | 5.69 | 0.23 | 15 |
| 019 | Discovery 600 | GE MEDICAL SYSTEMS | 4.65 | 0.22 | 5.68 | 0.17 | 15 |
| 114 | GEMINI TF TOF 64 | Philips Medical Systems | 6.94 | 0.13 | 6.37 | 0.13 | 15 |

**Supplementary Table 2**. SPITFIRE resolution estimator of ADNI FDG data.
